# Microbial Interactions from a New Perspective: Reinforcement Learning Reveals New Insights into Microbiome Evolution

**DOI:** 10.1101/2023.05.07.539711

**Authors:** Parsa Ghadermazi, Siu Hung Joshua Chan

## Abstract

Microbes are essential part of all ecosystems, influencing material flow and shaping their surroundings. Metabolic modeling has been a useful tool and provided tremendous insights into microbial community metabolism. However, current methods based on flux balance analysis (FBA) usually fail to predict metabolic and regulatory strategies that lead to long-term survival and stability especially in heterogenous communities. Here we introduce a novel reinforcement learning algorithm, Self-Playing Microbes in Dynamic FBA, that treats microbial metabolism as a decision-making process, allowing individual microbial agents to evolve by learning and adapting metabolic strategies for enhanced long-term fitness. This algorithm predicts what microbial flux regulation policies will stabilize in the dynamic ecosystem of interest in presence of other microbes with minimal reliance on predefined strategies. Throughout this article, we present several scenarios wherein our algorithm outperforms existing methods in reproducing outcomes, and we explore the biological significance of these predictions.

## Introduction

Microbes are present in almost all known biotic environments and their metabolism affects the flow of materials in their ecosystems. Microbes form intricate networks of interacting cells from various taxonomic branches with distinct functional traits which makes predicting their behavior challenging. However, determining the role of microbial life in their ecosystems can be a key to solving numerous challenges that we face today. Imbalance in human gut microbiome is consistently linked with diseases such as Inflammatory bowel disease [1]. At a larger scale, microbial metabolism is a major player in geochemical cycles on earth [2].

Metagenomics studies provides detailed information about the membership and biochemical functions of microbiomes. However, predicting the phenotype of microbial communities from their genotype is by nature a complex problem and has been an ongoing effort for the past few decades [3]–[6]. Trophic interactions between microbes is an important factor that significantly contributes to the evolution of microbiome composition and function in various ecosystems [7], [8] and it further complicates the prediction of emergent properties of microbial communities.

Understanding and predicting the dynamics of microbial systems has remained largely unknown despite the enormous growth in multi-omics techniques and it requires a wholistic modeling approach [9]. Mathematical models at different abstraction levels have been developed with the goal of making predictions that can explain the experimentally observed phenotypes [3], [6], [10]–[18]. GEnome-scale metabolic Models (GEMs) provide a detailed view of the biochemical networks of cells that are inferred from the genome of the organism of interest. GEMs generally contain up to thousands of biochemical reactions. Predicting the emergent properties of microbial communities by merely determining flux through such biochemical reactions is one of the main challenges in systems biology that yet remains to be addressed [5], [6]. Flux balance analysis (FBA) is a bottom-up approach that provides a scalable method for simulating cellular metabolism in the absence of reaction kinetic parameters [19]. FBA converts the system of differential equations resulting from mass balance across a cell to a linear programming problem by assuming steady state condition across the cells and defining a biologically relevant objective function[19]. Despite the defined objective function and the constraints on the flux values, FBA solutions are rarely unique, and feasible solutions form a large space where distinct phenotypes can coexist. Dynamic Flux Balance Analysis (DFBA) applies FBA in each timepoint, and using the calculated extracellular fluxes, changes in extracellular metabolites with time are calculated. These rates of changes are used in turn to form a system of differential equations that describe the concentration profiles of different species in the system over time [10], [20]–[29]. However, the problem of lack of a unique solution in FBA propagates through time. Consequently, in the cases where the attempt is to model the dynamics of a heterogenous microbial community, one is faced with an extremely open solution space where different solutions can represent significantly different phenotypes while all phenotypes can satisfy FBA requirements. More importantly, DFBA relies on instantaneous biomass maximization assumption. Although in simple cases this assumption might result in realistic simulations [10], in many other cases it fails to predict the observed behavior of microbial systems [12] because depending on the environment, maximizing instantaneous growth rate can result in low fitness in future or even extinction. For example, cells that excrete extracellular amylase to breakdown starch are spending energy to do so and lower their instantaneous fitness in turn. However, secreting amylase is required for degrading starch to smaller molecules such as glucose for the cells’ future use. Instantaneous biomass maximization will not allow any extracellular amylase secretion, unless previously set as a constraint on the model, while amylase secretion has been frequently observed in nature [30]–[32].

Therefore, it is important to put the concept of Nash equilibria and evolutionary stability in this context of metabolic interactions [9], [12]. A remedy proposed by Zomorrodi and Segre [33] was to determine Nash equilibria of the systems with several metabolic strategies of interest pre-defined. It beautifully captures the experimental results of some well-known microbial games of metabolic interactions. However, in general, it is virtually impossible to enumerate all possible metabolic strategies because of the high dimensional and continuous nature of the solution space defined by the mass balance constraints, directionality constraints, and nutrient availability. Therefore, an algorithm that can cover the entire possible solution space to a satisfactory extent when determining these stable interactions, and meanwhile does not rely on the instantaneous biomass maximization assumption, will greatly improve capability of predicting stable microbial interactions.

In this article we aim to address these challenges by introducing a new modeling approach that integrates deep reinforcement learning into DFBA to model microbial metabolism in a microbiome as a decision-making process. From this perspective, microbial cells evolve by trying different metabolic strategies and learning how to improve their long-term fitness by tuning their behavior using a reinforcement learning algorithm. In this framework, each GEM is modeled as an agent capable of making decisions. The decisions in this context are flux regulations in the metabolic network and the agents make these decisions using the observable environment states. Assuming that “bad decisions” are filtered through the natural selection process, we use reinforcement learning algorithms to find the strategies that lead to the long-term optimal behavior of microbes in the system that they are interacting with. In other words, microbial models learn how to interact by trial and error in their environment through self-play mechanism [34], without the need to pre-define metabolic and regulatory strategies. Reinforcement learning has shown great promise in solving very complex problems in the past decade [35]–[37] and have been used with success in different fields of science and engineering [38]–[43]. Although still relying on FBA, this approach is fundamentally different from biomass maximizing agents assumed commonly in traditional FBA and DFBA as the long-term consequences of actions are also considered in a dynamic context to find strategies that are also performing well in future rather than only an instance of time. In several cases, discussed shortly, greedily optimizing for biomass production will lead to early community extinction. Rationally, such strategies should be eliminated by the natural selection process. The strategies taken by RL agents after training can be useful to understand why certain types of behaviors are observed in real microbial systems.

## Methods

### Reinforcement Learning

In a reinforcement learning problem, one or more agents interact with their environment. An agent is an entity that is learning by interacting with the environment and is capable of *making decisions* in a given state. An environment can be defined as the collection of all entities that are surrounding the agent. Depending on the problem of interest, the agents can observe the entire or a part of their environment and based on a mapping called **policy** decide what actions to take in each state. Through this interaction with the environment the agents receive a reward and go to the next state according to the environment dynamics until it reaches the terminal state, that is when one **episode** is completed. The final goal of an agent is to maximize return during an episode. Return at time t is defined as the discounted sum of the collected reward after time t until the end of the episode [44]:

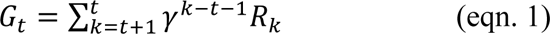

Here γ is the discounting factor which determines the importance of future rewards and *R*_t_ is the reward at time t.

Policy is a function that describes the behavior of an agent in each state. Policy function π is a mapping that outputs the probability of taking action *a*_t_ when the agent is in state *S*_t_ [45]:

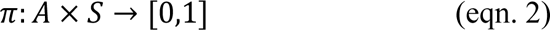

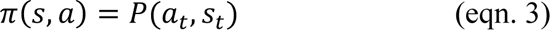

Another important definition is the **value function** under policy π which is defined as [44]:

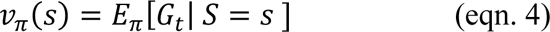

Value for state s under policy π is the expected return of being in state s and following policy π for the upcoming states [44]. Let’s consider two states *S*_1_, *S*_2_. If *v*_π_(*S*_1_) > *v*_π_(*S*_2_) then this means that if the agent is following policy π, the *expected* return is higher when the agent is in state *S*_1_ than when in *S*_2_. In other word the value function tells us how valuable it is to be in state s when following a specific policy function.

Depending on the type of problem at hand, there are families of RL algorithms that can be used [44], [46]. In the proposed framework (described in detail in ‘*<u>Self-PlAying Microbes in</u> <u>Dynamic Flux Balance Analysis (SPAM-DFBA)</u>*’), the states are observable extracellular metabolite concentrations which are continuous variables. On the other hand, actions are the reaction fluxes which are also continuous variables. This means that both the action and state space are continuous and multidimensional. For this type of problem, *policy gradient* family of algorithms is a good choice [44]. Among them, Proximal Policy Optimization (PPO) [47]is a policy gradient algorithm which has been used with success in many Artificial Intelligence problems recently [47]–[50] and is appropriate for modeling microbial communities.

### Proximal Policy Optimization

In policy gradient algorithms, the policy can be defined as a parameterized function, here a neural network, which maps a state to a real number, and the parameters of this function are tuned in a way so that the agent takes actions that lead to higher return because of the policy gradient theorem [51]. One issue with using reinforcement learning algorithms is that the underlying mathematical operation in our algorithm is a linear programming problem. The suggested actions by the policy function can easily form a solution that is not in the feasible region of the underlying LP problem and infeasible region can be in the proximity of areas where the return is maximum. This complicates the training process and we observed that many of the RL algorithms that we tested, failed on simple test cases. PPO addresses this issue by avoiding abrupt changes in the policy space. In the PPO algorithm, the agent tries to maximize the following surrogate objective function:

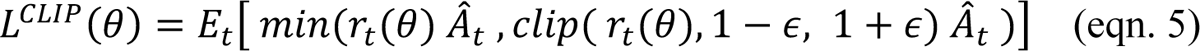

Where 

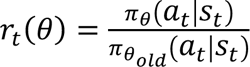

and

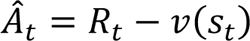

Setting this objective function (*L*^CLIP^) causes the policy function to increase the probability of actions that lead to higher return during an episode. In eqn. 5, *r*_t_(θ) represents the ratio of the probability of taking action *a*_t_ at state *S*_t_ under the improved and the old policy. *A*^^^_t_ is called the advantage function which is the difference between reward collected at state s and the expected return of states. The *clip* function does not allow for *r*_t_(θ) to go beyond 1 − ∈ and 1 + ∈. This objective definition has two implications on the policy function:

1. Probability of actions that result in more positive advantage will increase and the reverse will happen in the case of low advantage values.
2. The parameters of the policy function will only change if the change in the policy space is in a limited range.

Using this technique, PPO effectively stabilizes the training process. ε is a hyper parameter that depends on the problem of interest. In all of our simulations we used ε = 0.1.

The value function and the policy function (called critic and actor networks respectively) in all of our experiments are neural networks with 10 linear layers followed by hyperbolic tangent activation function [52] with Adam optimizer as the network optimizer in both cases. In all simulation cases we used 0.001 and 0.0001 as learning rates for critic and actor network respectively.

### Self-PlAying Microbes in Dynamic Flux Balance Analysis (SPAM-DFBA)

SPAM-DFBA is a dynamic algorithm. Every DFBA simulation happens over a defined period of time called episodes. When the final timepoint of the episode is reached, the agent go to the terminal state and a new episode is started. In the beginning of the algorithm the agents are defined by assigning them a metabolic model. At each time point, or “state”, the agents observe a part of their environment, by sensing the concentrations of some of the extracellular metabolites. According to their policy function, a neural network in our implementation, these agents regulate a subset of fluxes through their metabolic network in each state by posing constraints on the reaction flux bounds. Decisions on uptake fluxes are limited by the user-provided kinetic expressions and parameters. If the flux through an exchange reaction is higher than what is allowable by the corresponding kinetic rates, the flux value is clipped to the highest value that is allowable by the kinetic rates for that compound. The constraints imposed by the policy network are coming from feeding the state to the actor function. The main difference between SPAM-DFBA and the standard DFBA is that during FBA for an agent at each timepoint, additional constraints returned by the agent’s policy function are imposed. Next, FBA is performed, and the rewards, actions and states are recorded in an array until the end of the episode, Figure 2. Depending on the resources available, parallel episodes are simulated for the current policy at the same time. After one batch of episodes is finished, the critic network, the value function, gets updated based on the states and reward values. This step in reinforcement learning training process is called policy evaluation[45]. Next, according to the PPO objective function (eqn. 5) and using the information collected during the batch of episodes, the actor function is updated using gradient ascent to improve the policy, Figure 2. This step is called policy improvement. As a result of this iterative process, the agents become better at taking actions that maximize their own return by trial and error in the feasible solution space.

**Figure 1.**
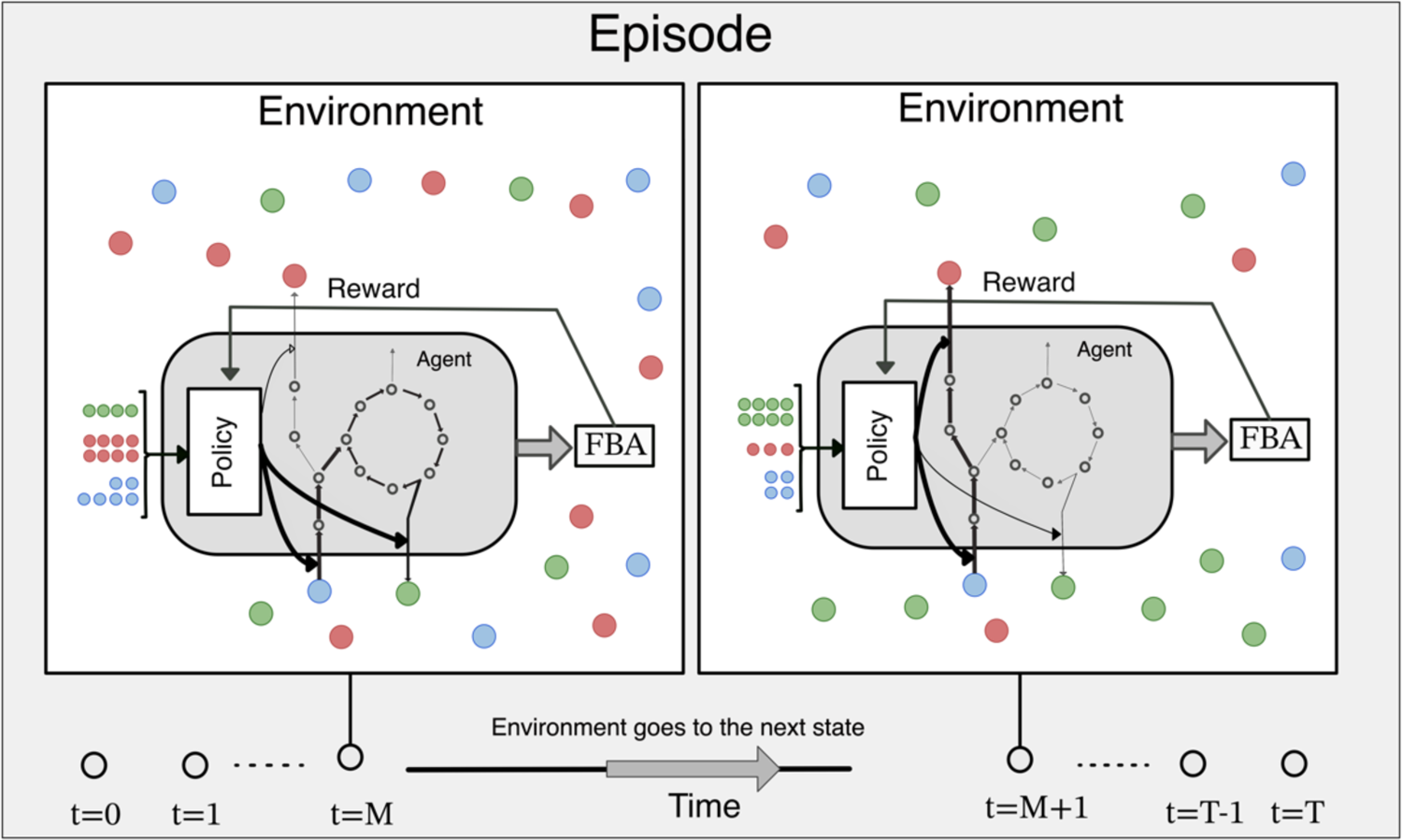
A schema of dynamic microbiome modeling viewed as a reinforcement learning problem. At each time step, the “agents” take an action according to their policy function without having a model of the environment dynamics and go to the next state. The actions in this sense are the constraints on the flux values. Based on the calculated flux with FBA a reward is given to the agent. Changes in the environment is calculated using the exchange fluxes and the environment goes to the next state. An episode contains all the time points from t=0 to t=T. After an episode is finished, the agents improve their policies to maximize the reward signals.

**Figure 2.**
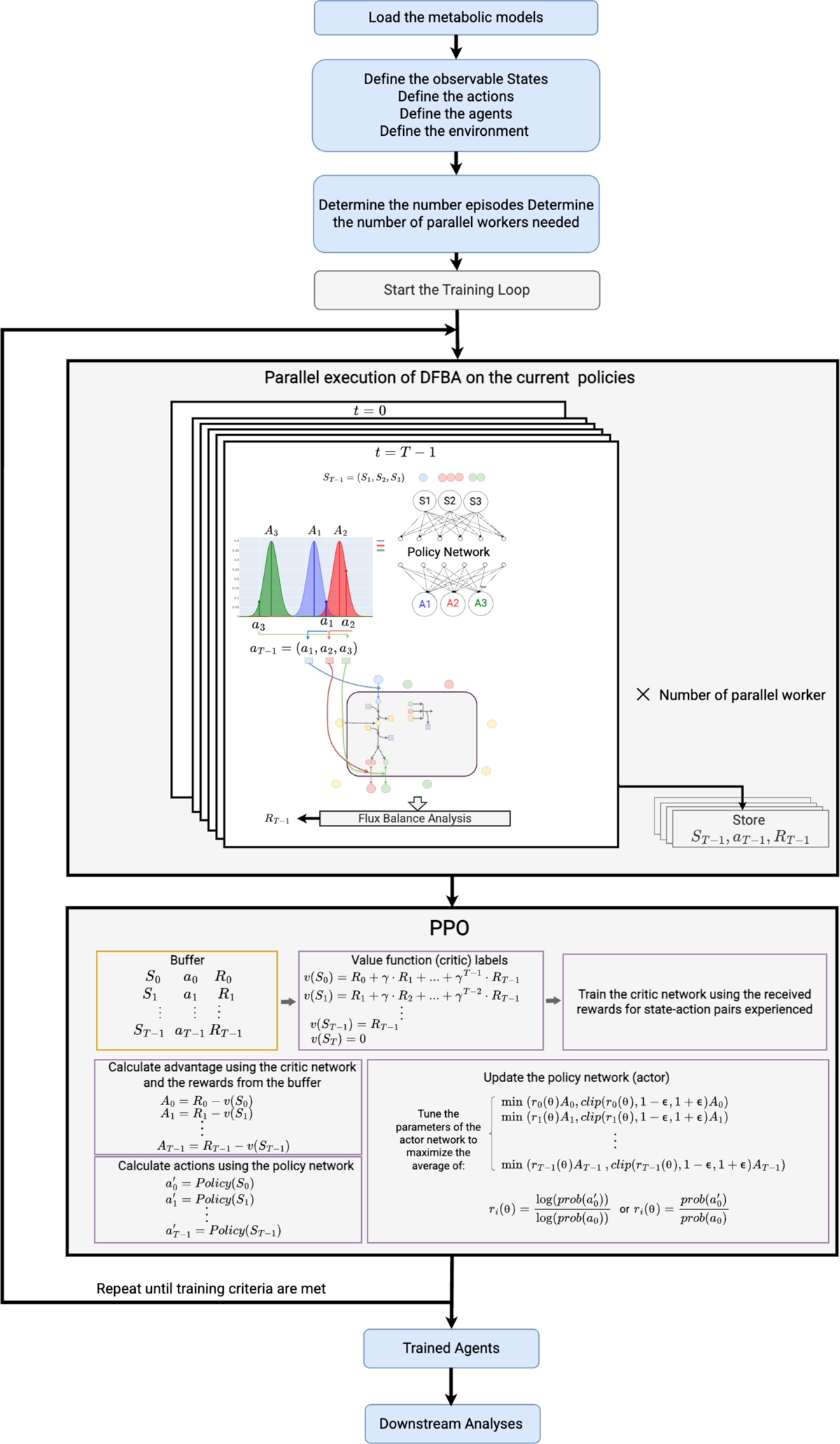
Schematic of SPAM-DFBA algorithm and a detailed view of using PPO in DFBA: Parallel workers run DFBA on the current policy for each agent as episodes of defined length, and collect the observations (States, rewards, and actions) made by the agent during each episode. Policy network of each agent suggest a set of actions: A1, A2, and A3. The agents take actions by drawing actual actions from a normal distribution around A1, A2, and A3 which constrains the lower bound of the corresponding reactions in the agent’s metabolic network. Afterwards, FBA is performed to calculate the flux for all reactions in the network. The collected observations enter a buffer that lasts for 1 batch of episodes. The information in this buffer is used to first update the critic network. Next, the policy network is updated in a way to maximize the PPO’s surrogate objective. The agents learn to act optimally by iterating this cycle until the policy improvement stops or other learning criteria are met.

In our implementation, the reward function has two components: a negative reward as penalty for infeasible flux distributions and a positive reward for growth/biomass production. When the policy network generates flux values that are outside of the feasibility range, the agent will get a negative reward to learn to stay away from infeasibility. For the positive reward for the growth rate determined by FBA at each time point, we emphasize that it is different from the maximization of biomass production for the immediate time point in those future rewards also affect the decision made by the agent in any timepoint. As a result, an action with low immediate reward but high future reward might be favored over immediate biomass optimization strategy.

### Implementation

SPAM-DFBA is implemented in Python. All the case studies are simulated with Python v3.10, and COBRApy v0.25.0 [53]. GLPK solver v0.4.7 was used to perform FBA in the toy communities and Gurobi optimizer [54] with academic license was used to perform FBA on the GEMs using COBRApy interface. PyTorch v1.12.0 [55] was used for building and training the neural networks. Same network structure and hyper parameters were used for all three cases to illustrate the robustness of this method with respect to the hyperparameters. Ray library was used to perform parallel computing [56]. Plotly v5.9.0 [57] was used to generate all the plots. Jupyter Notebooks are provided for reproducing all of the simulation cases. SPAM-DFBA is available as a package in PyPI with a detailed documentation website that facilitates using SPAM-DFBA for future studies. For more information, please refer to: https://github.com/chan-csu/SPAM-DFBA

## Results

We created multiple toy microbial communities that exemplify the weakness of FBA or DFBA which are inspired by NECom [12] to demonstrate the advantage of SPAM-DFBA. The following subsections will provide a detailed description of these biologically relevant toy communities.

## 1. Amylase secretion without mass transfer considerations

This group of toy communities were designed to emulate a case that microbial cells are grown on a mixture of starch and glucose in a well-mixed chemostat system, Figure 3A-C. The cells are capable of secreting amylase to degrade the available starch. However, producing amylase is an energy-consuming step in the organism’s metabolism and it requires ATP and precursors that would otherwise be used in biomass production. This poses a challenge on modeling the dynamics of such systems using DFBA because instant maximization of biomass would not allow any amylase secretion unless amylase production is set as a constraint on the underlying LP problem. Additionally, the amount of amylase secretion is also an important consideration as too much amylase production impedes growth in the environment. Exoenzyme production in microorganisms has been a system of interest in studying microbial games because of the cheater-producer coexistence problem [58]. Cheaters that do not secrete exoenzyme might evolve from an exoenzyme producer population because they benefit from the oligo-/mono-mers released by exoenzyme secreted by producers. In SPAM-DFBA, the agents take future awards into consideration in regulating their metabolic flux. As a result, they can learn to control amylase production in a manner that promotes their long-term survival by exploring various strategies in the environment and refining their policies depending on the rewards received.

**Figure 3.**
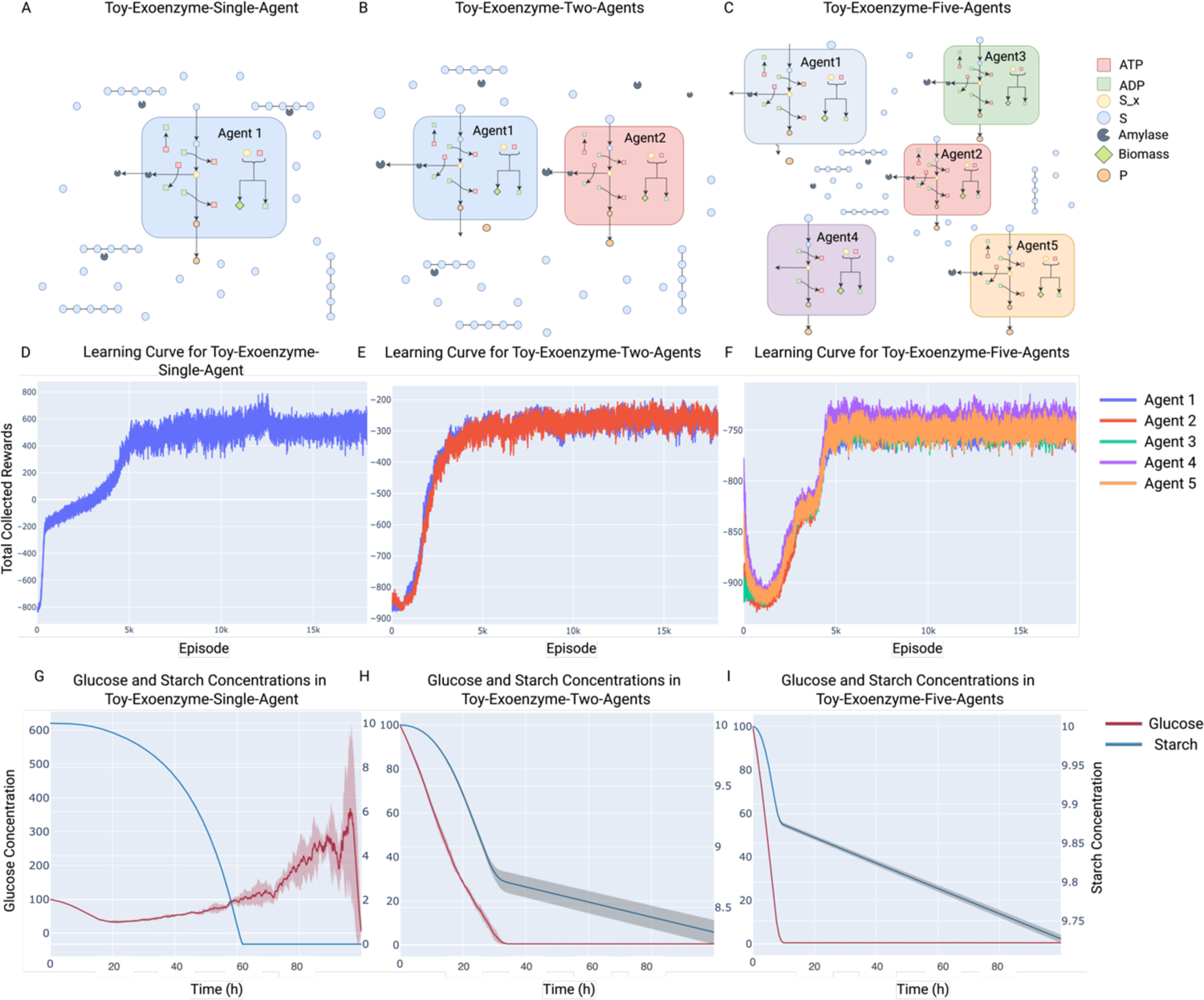
Training agents in a well-mixed chemostat system with starch and glucose initially present in the system. A-C) Schemas for the toy communities: Toy-Exoenzyme-Single-Agent, Toy-Exoenzyme-Two-Agents, and Toy-Exoenzyme-Five-Agents, respectively. D-F) Learning curve of the agents in Toy-Exoenzyme-Single-Agent, Toy-Exoenzyme-Two-Agents, and Toy-Exoenzyme-Five-Agents after training on 5000 batches of 4 episodes, respectively. Learning curves show the total collected rewards during an episode change over the course of the training process. Due to the energy required for maintenance, in multi-agent environments most agents receive negative rewards with the absence of sugar. G-I) Starch and Glucose concentration over time in Toy-Exoenzyme-Single-Agent, Toy-Exoenzyme-Two-Agents, and Toy-Exoenzyme-Five-Agents, respectively. The solid lines represent the mean value across all episodes in a batch and the shades represent 1 standard deviation across all episodes in a batch. Note that each actor acts randomly around the mean of the actor network output with standard deviation of 0.1. The left axis in each plot shows glucose concentrations and the right axes show the starch concentrations.

### 1.1 Convergence and amylase secretion predicted in single-agent simulations

We tested what strategies the intelligent agents in SPAM-DFBA will learn in terms of exoenzyme production when only one homogenous population exists (single-agent simulations, Figure 3A) vs. multiple phenotypes are allowed (two- and five-agent simulations, Figure 3B-C). In all cases, the agents converge to a stable policy which cannot be further improved (Figure 3D – F). There are significant growth and starch degradation in all cases (Figure 3G – I), suggesting the capability of the algorithm to identify feasible and biologically relevant solutions.

### 1.2 Agents learn to minimize exploitability in multi-agent environments

Simple DFBA cannot predict any amylase production by the metabolic network as it contradicts the biomass maximization assumption. However, SPAM-DFBA not only predicts amylase secretion on starch, but also reveals an interesting pattern when comparing the overall community growth and starch degradation between the single-agent and multi-agent cases. Starch utilization significantly decreases when the number of agents increases (Figure 3G – I). This trend suggests that when more agents are present in the environment, they become more conservative in terms of secreting amylase, granted spatial homogeneity. To look deeper into this observation, we examined the policy of the agents trained in different cases. The policy of the agent in Toy-Exoenzyme-Single-Agent is significantly different from the agents in Toy-Exoenzyme-Two-Agents (Figure 4A, B). When the agent is trained in Exoenzyme-Single-Agent, the glucose resulted from breaking down starch can be utilized only by the agent itself. However, when other agents exist in the environment this is not the case. Other agents can learn to cheat because of the randomness in their behavior and take up the available glucose without paying the cost for building the amylase molecules. As a result, in the single-agent environment amylase secretion is negatively correlated with the glucose level, the optimal policy instructs the agent to secrete amylase when the glucose level is low, so that the low glucose concentration is compensated by breaking down the available starch. On the other hand, secreting amylase when the glucose level is low in environments with more than one agent is risky. In this case, at low glucose level cheating can have a more deteriorative effect on the amylase producer organism. For this reason, in environments with more than one agent amylase production increases with glucose level.

**Figure 4.**
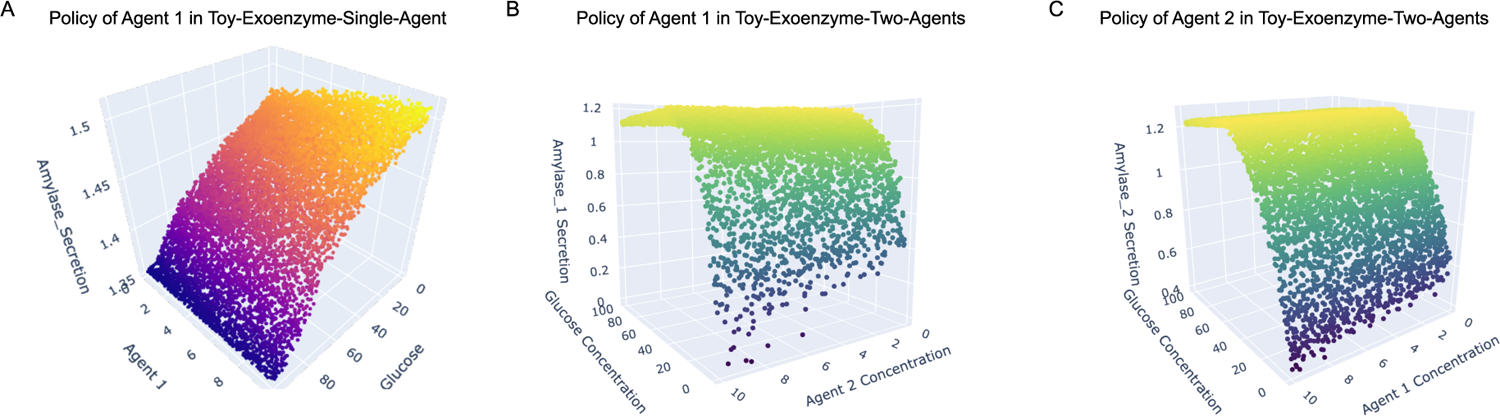
The policy profiles of agents in Toy-Exoenzyme-Single-Agent and Toy-Exoenzyme-Two-Agents environments after training on 5000 batches of 4 episodes. These plots are created by randomly generating 10000 points in the policy space of the trained agents. A) Policy of Agent 1 in Toy-Exoenzyme-Single-Agent with respect to glucose and Agent 1’s concentration B) Policy of Agent 1 in Toy-Exoenzyme-Two-Agents with respect to glucose and Agent 2’s concentration C) Policy of Agent 2 in Toy-Exoenzyme-Two-Agents with respect to glucose and Agent 1’s concentration.

What follows from this observation is that in a multi-agent environment, the agents that are trained in multi-agent environments should perform better than when trained in a single-agent environment as they have experienced different aspects of coexisting with other agents, such as cheating by other agents. To test this statement, we created a new two-agent environment, Toy-Exoenzyme-Single-Two-Comb. One agent is selected from Toy-Exoenzyme-Single-Agent and the other from Toy-Exoenzyme-Two-Agents. Supplementary Figure 1 shows that agent 1 that is trained in Toy-Exoenzyme-Single-Agent achieves significantly lower return compared to agent 2 that is trained in Toy-Exoenzyme-Two-Agents. Furthermore, the level of starch utilization is higher than Toy-Exoenzyme-Two-Agents but lower than Toy-Exoenzyme-Single-Agent which points to the fact that high amylase production by agent 1 is exploited by agent 2. This is an example of an emergent property of microbiomes which is made possible by considering the long-term effect of actions and explains the observed deterioration of performance in systems where large molecules are broken down by microbial cells.

## 2. Amylase secretion with mass transfer considerations

An important question is that how mass transfer rate in an environment can change this cheating behavior. Higher mass transfer increases the possibility of components moving away from the producers and be utilized by the cheaters. To put this hypothesis to test and see if our algorithm can predict lower mass transfer rate helps amylase producers, we simulated two five-agent environments. In one environment the mass transfer rate is lower than the other. Note that SPAM-DFBA does not consider spatial variations. However, we used a simple trick to simulate the effect of high and low mass transfer rate on the microbial strategies. More detailed information about this implementation can be found in the Jupyter Notebook titled “Case_Study_1_Starch_Amylase” in the documentation website and the GitHub repository for this project. The result of this experiment agreed with our hypothesis and the agents in the environment with lower mass transfer rate achieved higher return and higher starch utilization, i.e., lower final starch concentration, Supplementary Figure 2.

## 3. Toy-NECOM-Auxotrophs

Metabolite exchange between two auxotroph strains is another type of interaction that has been observed frequently in nature [59]. Modeling a community containing such strains with DFBA is problematic. In many cases, such as amino acid exchange, biomass maximization assumption in DFBA does not allow secretion of an amino acid. However, secretion of such compounds can be beneficial in the long run as the auxotrophs rely on the metabolic product of the other strains to survive. This problem becomes even more interesting since usually different strains of same species could compete for the same resources. To see if SPAM-DFBA can predict metabolite exchange between auxotrophs, Toy-NECOM-Auxotrophs environment was created. The schematic description of this environment is provided in Figure 5-A. Although they cannot survive on their own, the agents can grow by exchanging A and B. Each agent can sense its own concentration, the concentration of the other agent, concentration of S, concentration of extracellular A, and the concentration of extracellular B. Figure 5 shows the result of training the agents in this environment.

**Figure 5.**
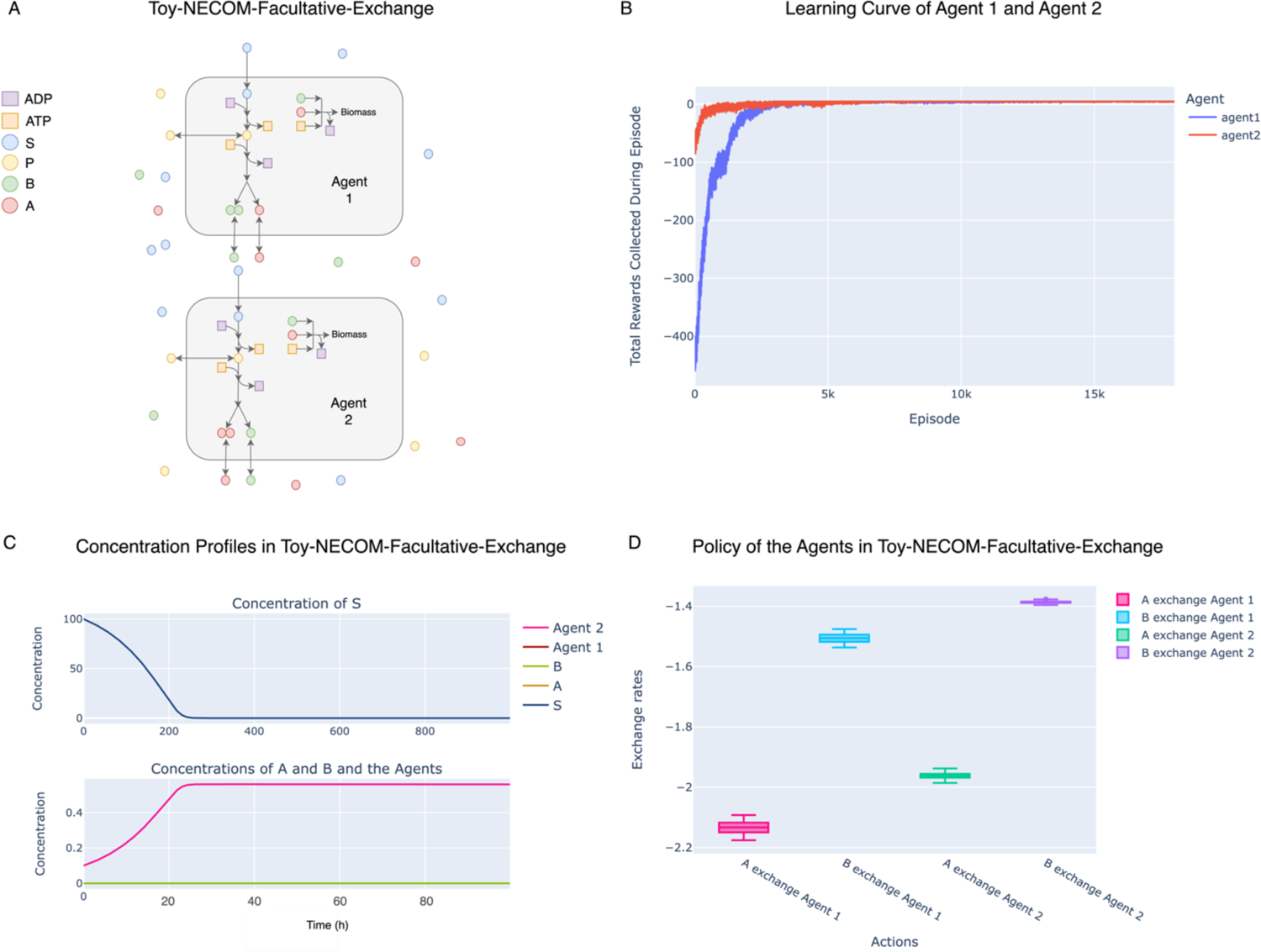
Training two agents in Toy-NECOM-Auxotrophs environment. A) A schematic view of the environment. In this case agent 1 and agent 2 have similar metabolic network with only one difference. Agent 1 cannot produce the biomass precursor A and Agent 2 cannot produce the precursor B. B) Learning curve of the two agents during 5000 batches of training. C) Concentration profile of the species in this environment over time in a batch of 4 episodes. D) Policies learned by the agents. Negative sign for fluxes means uptake and positive sign means secretion. Agent 1 learns to uptake whatever A that exists in the environment while secreting B for agent 2. Agent 2 has learned the opposite strategy which agrees with their mutation.

After training in this environment, the agents learn exchanging precursors A and B with each other. This prediction is qualitatively similar to the prediction by NECom [12].

### 3.1 Simulating adaptation in new environments

This community describes a scenario where two agents can synthesize both A and B but with different efficiencies. Supplementary Figure 3 shows the result of training the two agents in this community. Both agents learn to selfishly take up any A and B and not to cooperate which is in contrast with *<u>Toy-NECOM-Auxotroph</u>*. This is in contrast to the prediction of FBA when community biomass maximization assumption is used, cDFBA.

If metabolite exchange in this environment is dictated merely by the fact that cells rely on each other for survival, what happens if we supplement the Toy-NECOM-auxotroph environment agents with A and B externally after training? The underlying hypothesis is that this richer environment should discourage the cooperation between the two complementary auxotrophs. To answer this question, we created a new environment, Toy-NECOM-Auxotrophs-Shift. We used the two auxotroph agents from Toy-NECOM-Auxotrophs and simulated a scenario where A and B initially is supplemented in the environment. This case is designed to predict how changes in environment can shape the trophic behavior of microbial communities.

Figure 6 shows that the auxotrophic agents shift their policy from metabolic exchange to selfishly taking up A and B (negative median flux) when A and B is supplemented externally. This prediction is consistent with previous experimental results that supplementation of the needed metabolites discourages the cross-feeding interactions between auxotrophic mutants [60]. This also shows an intriguing capability of SPAM-DFBA to predict how a certain microbial population adapted to an environment might evolve in a new environment.

**Figure 6.**
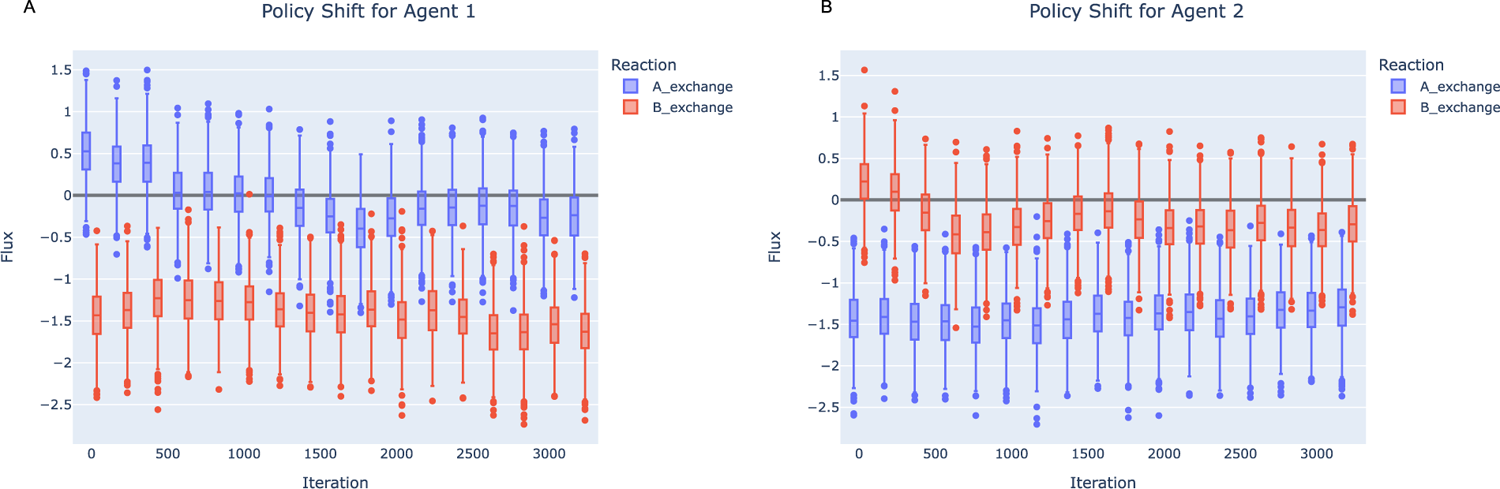
Policy shift after 3000 iteration of training agents of Toy-NECOM-Auxotrophs environment in Toy-NECOM-Auxotrophs-Shift. Both agents shift their policy from cross-feeding to taking up both A and B as much as possible. Here, the box plots show the range of actions across randomly generated states to represent the policy function.

## 4. E. coli Auxotrophs

So far, we only discussed small toy examples. IJO1366-Tyr-Phe-Auxotrophs environment was created to prove the scalability of our approach to a community of genome-scale models, IJO1366 for *E. coli* K-12 MG1655 [61]. In this environment two *E. coli* auxotrophs were made. One mutant could not synthesize tyrosine and the other could not synthesize phenylalanine. Neither of the mutants could grow alone on M9 minimal medium. However, after training the agents converged to a policy that they would exchange the amino acid that they can produce and grow by using the amino acid secreted by the other agent. In this environment phenylalanine mutant grew significantly more than the tyrosine mutant (Figure 7). This observation is experimentally validated for the same system by Mee, et al. (2014) [59] where the exchange of amino acids between the strains and the prevalence of the phenylalanine mutant has been verified (Agent 2 in our simulation). No phenotypic data from this or other experiments was used during the training process, which shows the promise of the assumption that evolution favors traits at individual level that leads to high long-term fitness of the cells.

**Figure 7.**
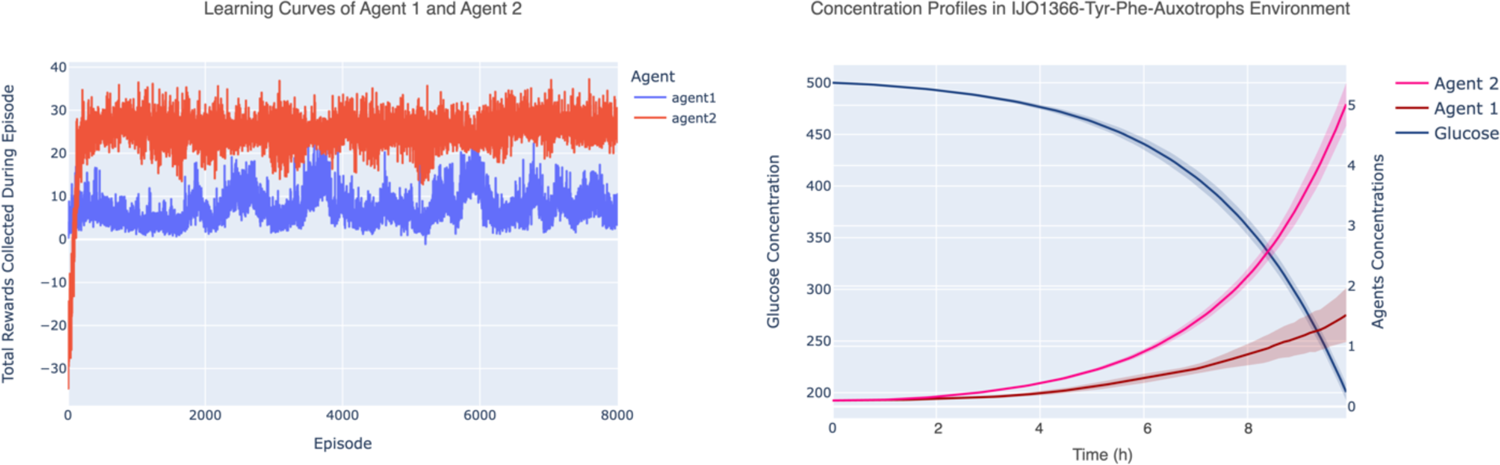
Training E. coli mutants in IJO1366-Tyr-Phe-Auxotrophs environment. Agent 1 is Tyrosine mutant and Agent 2 is Phenylalanine mutant. A) Learning curve of Agent 1 and Agent 2. Agent 2 receives higher return than Agent 1 although it starts from worse policy. B) Concentration profiles for glucose and the agents. It is also obvious from B that Agent 2, phenylalanine, achieves superior growth compared to Agent 1, tyrosine agent.

## Discussion and Conclusion

In this article we presented a novel algorithm that provides insights into microbial interactions by allowing the microbial agents to freely explore flux regulation strategies and select metabolism regulation strategies that lead to their higher long-term fitness of the agents. This way we can explain the observed phenotypes in multiple communities that current algorithms fail to explain. Defining this problem in a FBA framework forces the strategies to be inside a space where mass balance and flux constraints are still satisfied. Another advantage of using FBA is that the underlying LP problems can be solved efficiently. We examined this algorithm on multiple test scenarios that emulate biologically relevant scenarios.

In scenarios where agents should coexist with other agents, the agents learned aspects of interacting with others while still tried to maximize their own return. The outcome of starch-amylase system has an interesting interpretation. Cheating in microbial communities can significantly affect the amount which large molecules such as starch are degraded in hydrolysis [50], [62]–[72]. Taking spatial heterogeneity into consideration revealed that in communities with higher mass transfer limits, the agents secrete more amylase and starch utilization becomes higher (Supplementary Figure 2). The reason behind this observation is that low mass transfer implies that the glucose that is produced by an agent will stay away from the other agent that could possibly cheat, and in turn, the agents will see more positive signal by secreting amylase.

With this algorithm we were able to explore other types of microbial interactions in a dynamic context. One problem that we were interested in was that whether we can explain metabolite exchange between auxotrophic strains through this framework [59], [73]. We hypothesized that without any predetermined exchange strategies or community level objectives, the optimal agents can find metabolite exchange with other agents strategy to maximize their own long-term fitness. Optimal agents in Toy-NECOM-Auxotrophs learned that exchanging A and B will increase their long-term fitness.

To see if this algorithm can be used for genome-scale model in real environments, we created an environment of two E. coli auxotrophs, tyrosine and phenylalanine, inspired by the experiments in [59]. Although we did not use any sort of experimentally observed phenotypic data, the agents learned to exchange the amino acid that they can produce, and the other agent cannot. Interestingly, our simulations indicate that the phenylalanine mutant achieves superior growth compared to the tyrosine mutant, Figure 7, which follows the same trend as is experimentally observed and reported in [59]. Being able to predict such emergent behaviors of microbiomes by purely relying on metabolic capability of the cells and ecological first principles is what distinguishes SPAM-DFBA from the other existing algorithms.

An interesting study [60] reported the behavior change of auxotrophs when inserted in an environment that supplies all the components that they need for growth. In this scenario they shift their exchange strategy to uptake all the compounds from the environment and stop secreting the metabolites further. Our simulations showed similar shift for auxotrophic agents which reflects that the agents adapt their strategies according to the changes in the environment, Figure 6, and shows assuming that cells are *maximizing their own long-term fitness* can reproduce several real scenarios is missed by simple DFBA.

Previous cases revealed that agents that depend on each other for survival will evolve to exchange metabolites with each other and when this strict dependence doesn’t exist anymore selfish behaviors emerge. Toy-NECOM-Facultative-Exchange provides more evidence for this trend. In this case, if a community level objective such as, total community biomass maximization, is defined then there will be A and B exchange between agent 1 and agent 2 [12], [74]. This is the result predicted by the direct extension of FBA where a microbial community is optimized as one compartmentalized model. However, this is not what SPAM-DFBA predicts. In this case, A and B exchange strategy is exploitable by the agents. Since the agents do not rely on each other for survival, any exchange of A and B is exploitable by either of the agents in the case of resource limitation. Consequently, the agents finally adhere to taking up any A and B, limited by the kinetic rules provided for the model, that exist in the environment which is shown in Supplementary Figure 3. This is consistent with the previous NECom prediction and game-theoretical analysis [12], and exactly matches simple DFBA prediction.

SPAM-DFBA is well suited for answering important questions in the field of microbiology by predicting the emergent behavior of microbiomes using metabolic capability of the cells in contrast with commonly used ecological models such as Generalized Lotka-Volterra [75]–[77], which do not base their predictions on the metabolic network of the microbes. SPAM-DFBA is a dynamic framework, and the environment changes such as resources limitations can be simulated while methods based on FBA, and not DFBA, cannot make such considerations which is critical and can significantly shape microbial interactions [78].

Another advantage of this approach is that optimization is done at individual model level instead of community level objectives. This means that unrealistic interactions discussed in detail in [12] are avoided. If a particular random action is advantageous to the long-term fitness of an agent, this behavior gets reinforced in the policy of the agent using the PPO algorithm. We believe that this has a lot of similarities to the process of natural selection. We would like to emphasize that our method does not imply that the microbial cells are intelligently seeking the optimal behavior in their environment. However, it is the resemblance of this algorithm to the process of natural selection that results in more realistic predictions for a given environment.

There are multiple ways in which SPAM-DFBA could be further improved in future studies. The current implementation can scale to multiple GEMs in a manageable amount of time. However, if simulations for a real microbiome with hundreds of taxa is desired this approach becomes intractable. More efficient implementations can help in this regard. One particularly interesting venue is to completely remove the need for LP solvers by letting the agents sample the feasible action space. This not only can improve the speed dramatically, but it also relaxes any assumption imposed on the metabolism of the agents under optimization formulation. Improvements in sample efficiency of RL algorithms can also improve the efficiency of this algorithm in future.

Hyperparameters such as clipping threshold or learning rates also affect the efficiency and stability of the learning process for the agents. Although we used same hyperparameters for all the case studies, optimal combination of the hyperparameters can be explored methodically either by exhaustive search or using appropriate optimization techniques [79], [80]. As an example, we have examined the effect of important hyperparameters on the learning process for “Toy-Exoenzyme-Single-Agent” which is included in Supplementary Figure 4.

In this article we just showed the potentials of formulating DFBA as a RL problem and discussed how this approach can predict microbial interactions in simple communities. Applying this approach to more complex ecosystems and validation with experimentally observed phenotypes is a natural next step for future studies.

## Data Availability

All of the source codes for recreating the simulations, data analysis, and generating figures are provided in the GitHub repository for this project: https://github.com/chan-csu/SPAM-DFBA

