## Supplementary Files for "Microbial Interactions from a New Perspective: Reinforcement Learning Reveals New Insights into Microbiome Evolution"

### Supplementary Figures

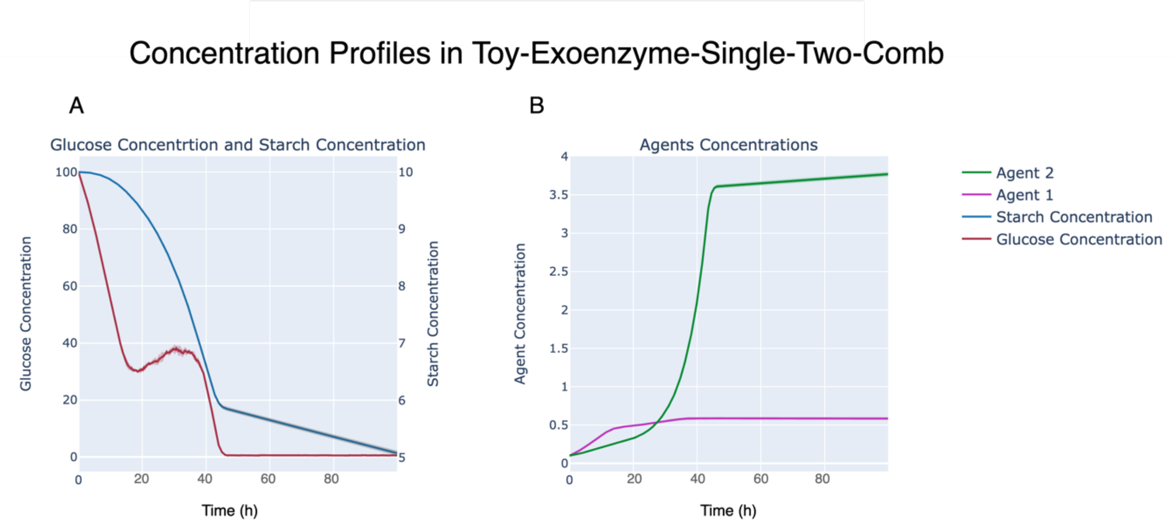


***Supplementary Figure 1*** Concentration profiles in Toy-Exoenzyme-Single-Two-Comb environment. A) Concentration of glucose and starch over time. B) Concentration of Agent 1 and Agent 2 over time. Agent 1 is trained in Toy-Exoenzyme-Single-Agent and Agent 2 is trained in Toy-Exoenzyme-Two-Agents.

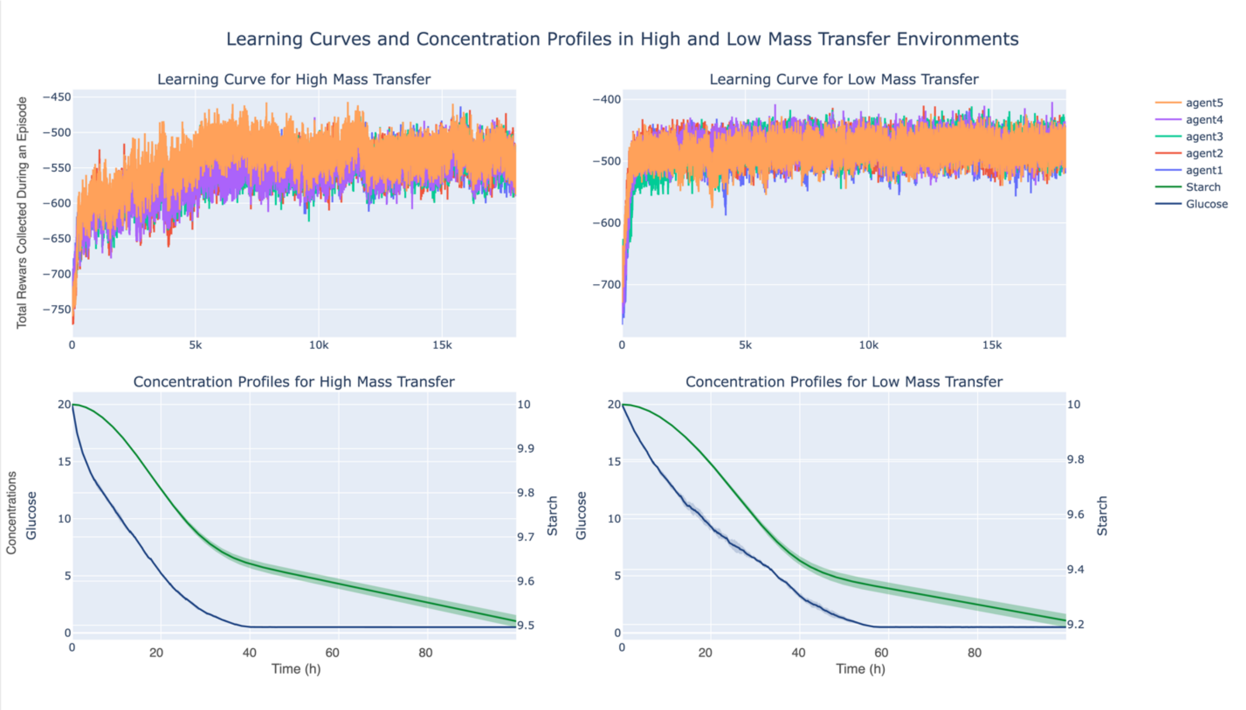


***Supplementary Figure 2*** The difference in the dynamics of starch degradation in two environments with high and low mass transfer rates. In the low mass transfer rate environment, the trained agents achieve higher returns, and degrade more starch when compared to the high mass transfer rate environment. This reflects the effect of cheating in microbial communities.

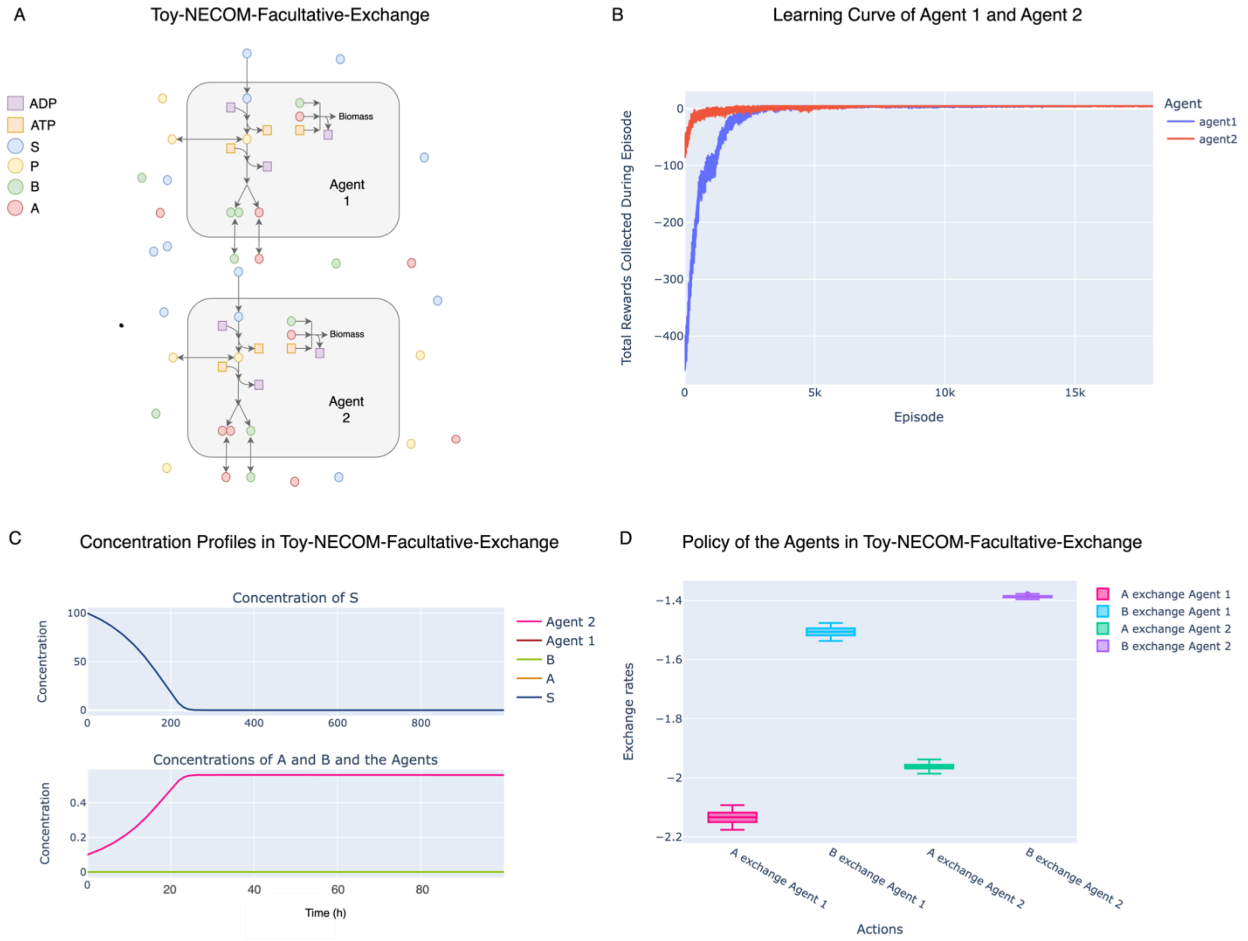


***Supplementary Figure 3*** Training two agents in Toy-NECOM-Facultative-Exchange environment. A) A schematic view of the environment. B) Learning curve of the two agents during 5000 batches of training. C) Concentration profiles after training for 5000 batches of 4 episodes. In this case the concentration profiles for extracellular A and B are exactly the same. Same is true for the concentrations of Agent 2 and Agent 1 D) The policies learned by the agents. Negative sign means uptake. Both agents learn to avoid any exchange of A or B and selfishly take up any A or B that is present in this environment. Note that if the absolute value of the flux for a reaction is larger than what the uptake kinetics allow, then the flux value gets clipped to match the highest value that is kinetically possible, which is the case for all exchange reactions here. This case yields exactly same results as DFBA.

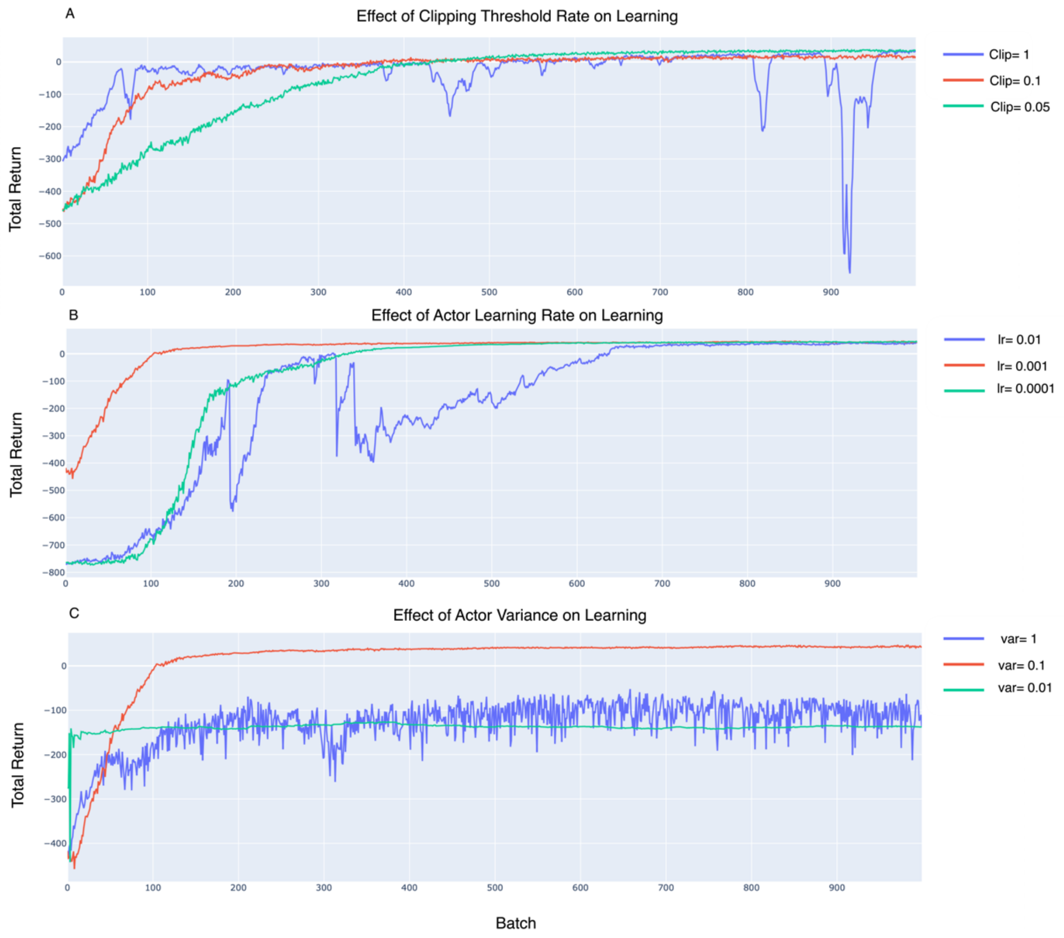


***Supplementary Figure 4*** Effect of different hyperparameters on learning performance in Toy-Exoenzyme-Single-Agent environment (1000 batches). A) choosing high clipping value will cause destabilization of the training process. This is happening because the policy is prone to higher shifts when the clipping threshold is high for the actor network output. B) Lower learning rate is more stable than high learning rates for similar reason in A. C) Very high and very low actor variance has a negative effect on the training process: High variance will result in higher chance of big jumps and more infeasible actions and sub optimal behavior. Low variance will result in less noise but also lower exploration and sticking in sub optimal policy.

**Supplementary Table 1** Characteristics of the tested environment. The durations are reported for Apple MacBook Air 2021 with M1 processor and 16 GB of RAM. For all simulations there are 4 episodes in each batch and 5000 batch per each simulation.

| Environment | Number of Agents | Number of Reactions (per agent) | Number of Metabolites (per agent) | Number of Time Steps per Episode | LP Optimization Time (s) (per agent) | DFBA Step Time (s) | Batch Time (s) | Simulation Time (s) |
|---|---|---|---|---|---|---|---|---|
| Toy-Exoenzyme-Single-agent | 1 | 9 | 7 | 1000 | 0.0003498 | 0.0005561 | 1.192 | 6135 |
| Toy-Exoenzyme-Two-agents | 2 | 9 | 7 | 1000 | 0.0004752 | 0.001435 | 3.011 | 15380 |
| Toy-Exoenzyme-Five-agents | 5 | 9 | 7 | 1000 | 0.0005903 | 0.003936 | 8.383 | 42710 |
| Toy-NECOM-Auxotrophs | 2 | 9 | 7 | 1000 | 0.0003250 | 0.0006016 | 1.058 | 6023 |
| IJO1366-Tyr-Phe-Auxotrophs | 2 | 2582 | 1805 | 250 | 0.02076 | 0.05958 | 16.75 | 84120 |
